# Neuroepithelial depletion schedules cessation of neurogenesis in the *Drosophila* optic lobes

**DOI:** 10.1101/2023.12.14.571634

**Authors:** Phuong-Khanh Nguyen, Louise Y Cheng

**Affiliations:** Peter MacCallum Cancer Centre, Melbourne, VIC 3000, Australia; Sir Peter MacCallum Department of Oncology, The University of Melbourne, VIC 3010, Australia; Department of Anatomy and Physiology, The University of Melbourne, VIC 3010, Australia

**Keywords:** *Drosophila*, medulla neuroblast, gliogenesis, neuroepithelium

## Abstract

The brain is consisted of diverse neurons arising from a limited number of neural stem cells. *Drosophila* neural stem cells called neuroblasts (NBs) produces specific neural lineages of various lineage sizes depending on their location in the brain. In the *Drosophila* visual processing centre – the optic lobes (OLs), medulla NBs derived from the neuroepithelium (NE) give rise to neurons and glia cells of the medulla cortex. The timing and the mechanisms responsible for the cessation of medulla NBs are so far not known. In this study, we show that the termination of medulla NBs during pupal development is determined by the exhaustion of the NE stem cell pool. Altering NE-NB transition during larval neurogenesis disrupts the timely termination of medulla NBs. Medulla NBs terminate neurogenesis via a combination of cell death, terminal symmetric division, and a switch to gliogenesis. We show that temporal progression is not required for the termination of medulla NBs. The timing of NB cessation can be altered through the acquisition of a glial cell fate via Glial cells missing, or through conversion to type II NB cell fate via Tailless, or by inhibition of differentiation via Prospero knockdown. As the *Drosophila* OL shares a similar mode of division with mammalian neurogenesis, determining how and when these progenitors cease proliferation during development can have important implications for mammalian brain size determination and regulation of its overall function.

## Introduction

The central nervous system (CNS) is the cognitive control centre of the body and is generated by neural stem cells (NSCs) during development. *Drosophila* NSCs, called neuroblasts (NBs), produce different neural and glial types and subtypes that contribute to the formation of the adult brain. The cellular diversity of the CNS is determined in a region-specific manner, via differential control of NB division mode and temporal identity <u>(reviewed by Harding & White, 2018)</u>. The majority of NBs are Type I NBs located in the ventral nerve cord (VNC) and the central brain (CB). They divide asymmetrically to generate a NB and a smaller ganglion mother cell (GMC), that undergoes terminal differentiation to give rise to neurons or glial cells <u>(Harding & White, 2018)</u>. In addition, there are eight Type II NBs located on the dorsal surface of the CB, where NBs asymmetrically divide to give rise to a NB and an intermediate neural progenitor (INP) <u>(Bello et al., 2008; Boone & Doe, 2008; Bowman et al., 2008</u>). INPs undergo maturation prior to additional rounds of asymmetric division, to give rise to neurons and glial cells.

The optic lobes (OLs) of the brain are the visual processing centre of the adult fly brain (Figure 1A). During neural development, neurogenesis in the OLs takes place in two proliferation centres: the inner and the outer proliferation centres, which produce different neural cell types that populate the various cortexes of the visual system (Sato *et al*, 2019; <u>Apitz & Salecker, 2014)</u>. In the outer proliferation centre (OPC), medulla NBs are generated from a pseudostratified neuroepithelium through a differentiating “proneural wave” (Figure 1A’) <u>(Egger et al., 2007; Li et al., 2013; Yasugi et al., 2008)</u>. This proneural wave is characterised by the expression of proneural factors such as Lethal of scute (L’sc) which precedes NB generation from NE cells <u>(Yasugi et al., 2008)</u>. During early larval stages, OPC NE cells undergo symmetric cell divisions to expand the progenitor population. By early-third instar, progression of the proneural wave is initiated in the NE cells at the medial edge, leading to their differentiation into medulla NBs. Medulla NBs sequentially express a series of temporal transcription factors <u>(Konstantinides et al., 2022; Li et al., 2013; Zhu et al.</u>, 2022). During the early-mid temporal stages, NBs divide asymmetrically to produce two neurons, upon acquiring the terminal temporal identity, NBs switch to become glioblasts that produce glial cells (Zhu *et al*, 2022).

**Figure 1:**
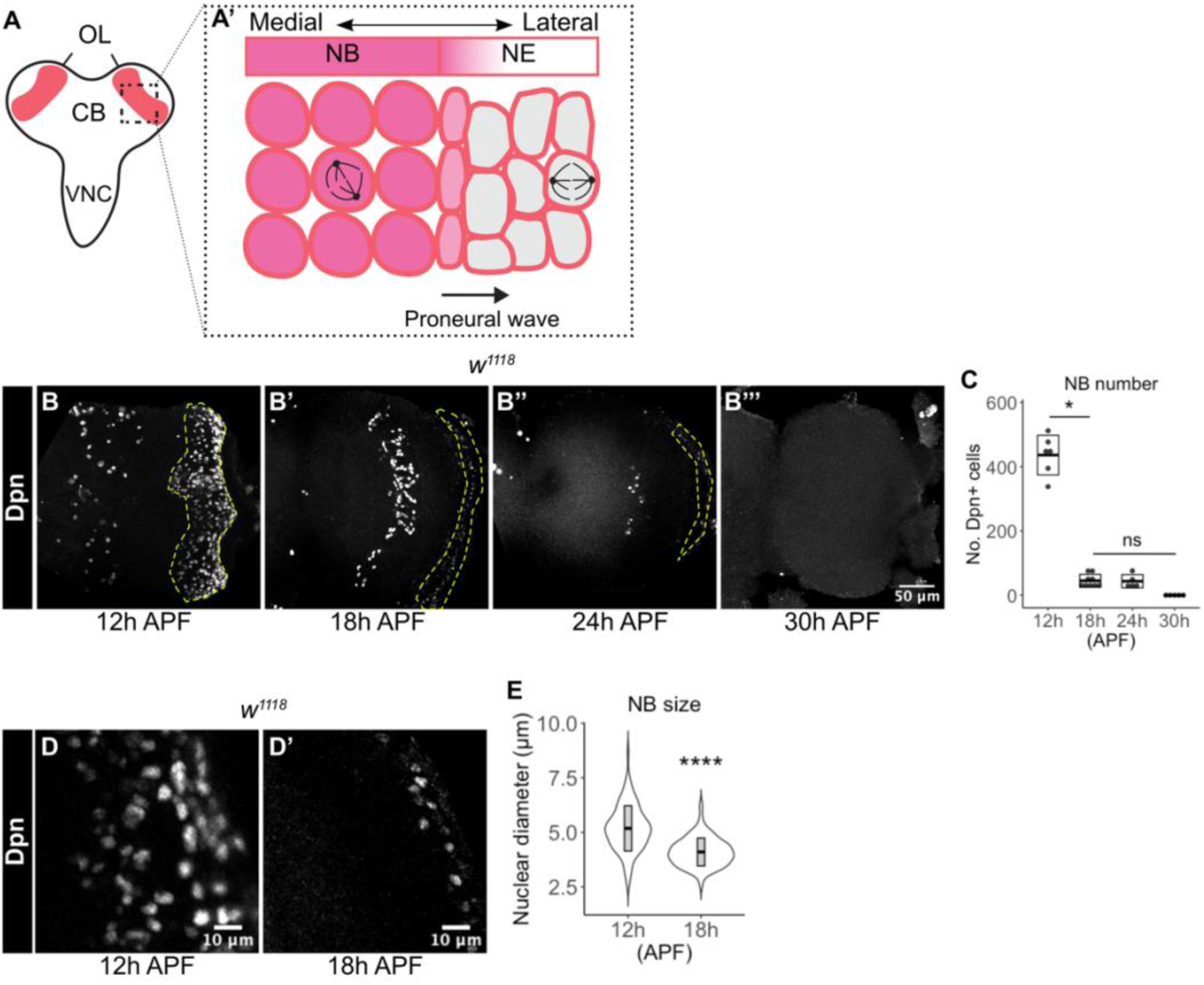
Medulla NBs are terminated during early pupal development. **(A-A’)** Schematic depicting the larval CNS that is consisted of three main regions: the central brain (CB), the ventral nerve cord (VNC) and the optic lobe (OL). Box area indicates inset shown in A’. (A’) The apical view of the outer proliferation centre (OPC) of the OL. Here, NE cells first symmetrically divide to self-renew. At mid-larval development, a proneural wave is initiated and travels from the medial to lateral of the NE. Upon the proneural wave, NE cells differentiate into NBs and switch to asymmetric cell division for neurogenesis. **(B-B’’’)** In the wildtype (*w^1118^*) OL, the number of medulla NBs marked by Dpn (dashed lines), gradually reduces from 12 hr to 24 hr APF. Medulla NBs completely disappear at 30 hr APF. **(C)** Quantification of the numbers of medulla NBs at 12 hr, 18 hr, 24 hr, 30 hr APF in the wildtype OLs. Kruskal-Wallis test and Dunn’s test to correct for multiple comparisons: *p = 0.0314. 12 hr: n = 6, m = 436.3 ± 25.21. 18 hr: n = 9, m = 45.22 ± 6.616. 24 hr: n = 5, m = 43.6 ± 9.255. 30 hr: n = 5, m = 0.000 ± 0.000. **(D-D’)** In the wildtype, the nuclear size of medulla NBs marked by Dpn decreases from 12 hr to 18 hr APF. **(E)** Quantification of the nuclear diameters of medulla NBs at 12 hr and 18 hr APF. Mann-Whitney test: ****p < 0.0001. 12 hr: n = 109 cells, m = 5.182 ± 0.099 μm. 18 hr APF: n = 61 cells, m = 4.100 ± 0.082 μm.

At the end of neurogenesis, NBs of different neural lineages are eliminated through distinct mechanisms. For instance, type I NBs in the VNC and the CB exit the cell cycle and terminally differentiate at around 24 hr after pupal formation (APF), whereas type I abdominal NBs are eliminated via apoptosis prior to pupation (Maurange *et al*, 2008; Bello *et al*, 2003; Ito & Hotta, 1992). Mushroom body NBs are the last NBs to undergo termination, and they do so through apoptosis or autophagy prior to adult eclosion (Siegrist *et al*, 2010; Pahl *et al*, 2019). So far, the timing and mode by which medulla NBs terminate is not known; furthermore, it is unclear whether the temporal series, which schedules NB cessation in other NB lineages <u>(Maurange et al., 2008; Syed et al., 2017)</u> is also involved in the termination of medulla NBs.

Here, we find that medulla NBs terminate between 12-18 hr APF. Unlike other NB lineages, the timing of medulla NB cessation is mainly regulated by the depletion of the NE pool. The exhaustion of the NE during pupal development results in the lack of the formation of new NBs. Hence, altered NE-NB transition is sufficient to change the timing of NB termination. Additionally, we demonstrate that medulla NBs terminate via a combination of cell death, size-symmetric terminal differentiation and a switch to gliogenesis. While medulla NBs undergo temporal transitions, the temporal series is not required for NB termination. However, overexpression of Tll, Gcm and knockdown of Pros prevents timely NB cessation during the pupal stages.

## Results

### Medulla NBs terminate during early pupal development

To characterize the timing at which medulla NBs are eliminated during pupal development, we first assessed the number of NBs in the OPC using the pan-NB marker Deadpan (Dpn). We found that the number of NBs was significantly reduced between 12-18 hr APF and by 30 hr APF, all the NBs were eliminated (Figure 1B-C).

In most regions of the CNS, NB termination is associated with a reduction in cell growth and proliferation rate, scheduled by a peak of ecdysone between larval and pupal development (Homem *et al*, 2014). To test if ecdysone signalling plays a role in the termination of medulla NBs, we first looked at the expression of the ecdysone receptor (EcR) in the medulla NBs at 12 hr APF, prior to the reduction in the number of medulla NBs in the OL. Immunostainings showed that EcR is expressed at very low levels in the medulla NBs at 12 hr APF (Supplementary Figure 1A). Additionally, when EcR was knocked down via RNAi in the medulla NBs with *eyR16F10-GAL4,* which is expressed in a subset of medulla NBs and their progeny <u>(Veen *et al*, 2023)</u>, we could not detect any persistent NBs at 24 hr APF (Supplementary Figure 1B, C, E). A similar phenotype was observed using a dominant negative form of EcR (EcR^DN^) (Supplementary Figure 1D, E). Together, these data suggest that ecdysone signalling in the medulla NBs is not required for their cessation during pupal development.

During larval development, NBs regrow to their original size after each division empowered by aerobic glycolysis providing the NBs with energy and metabolites for rapid cell growth and proliferation (Homem *et al*, 2014; Maurange *et al*, 2008). To assess if a reduction in cell size also precedes NB elimination in the medulla, we monitored the nuclear size of NBs marked by Dpn. Between 12-18 hr APF when the majority of medulla NBs disappear, we found that there was a significant reduction in the nuclear sizes of the medulla NBs (Figure 1D-E), suggesting that decreased cell growth precedes the cessation of medulla NB. However, increasing cellular growth through the overexpression of a constitutive active form of the PI3K catalytic subunit Dp110 (dp110^CAAX^) <u>(Leevers et al., 1996)</u>, or the activation of the cellular growth regulator Myc (Rust *et al*, 2018) did not prolong NB persistence in the OL at 24 hr APF (Supplementary Figure 1F, G, J, data not shown).

In type I NBs of the CB and VNC, it was previously shown that NB termination is dependent on core components of the mediator complex, which promote key enzymes to increase oxidative phosphorylation (OxPhos) to slow down NB regrowth (Homem *et al*, 2014). Nevertheless, neither inhibition of OxPhos itself via the knockdown of Complex I (Ameele & Brand, 2019), nor the knockdown of Med27 via RNAi (Homem *et al*, 2014), a component of the Mediator complex, significantly affected medulla NB persistence at 24 hr APF (Supplementary Figure 1H-J). As such, although cell size reduction coincides with NB elimination during pupal development, a decrease in cell growth, OxPhos, and the ecdysone-induced Mediator complex are not required for the termination of medulla NBs.

### The depletion of the NE determines the timing of medulla NB cessation

NBs of the medulla are continuously generated from NE cells from mid-late larval development by the proneural wave <u>(Egger et al., 2007; Yasugi et al., 2008</u>). To assess if NE depletion during pupal stages is linked to the cessation of medulla NBs, we first characterised the volume of NE cells during pupal development. From 12 hr to 24 hr APF, the NE gradually decreased in volume, concordant with the reduction in the number of medulla NBs during this time window (Figure 2A-B, Figure 1B-C). This suggests that the depletion of the progenitor NE pool may drive the cessation of medulla NBs.

**Figure 2:**
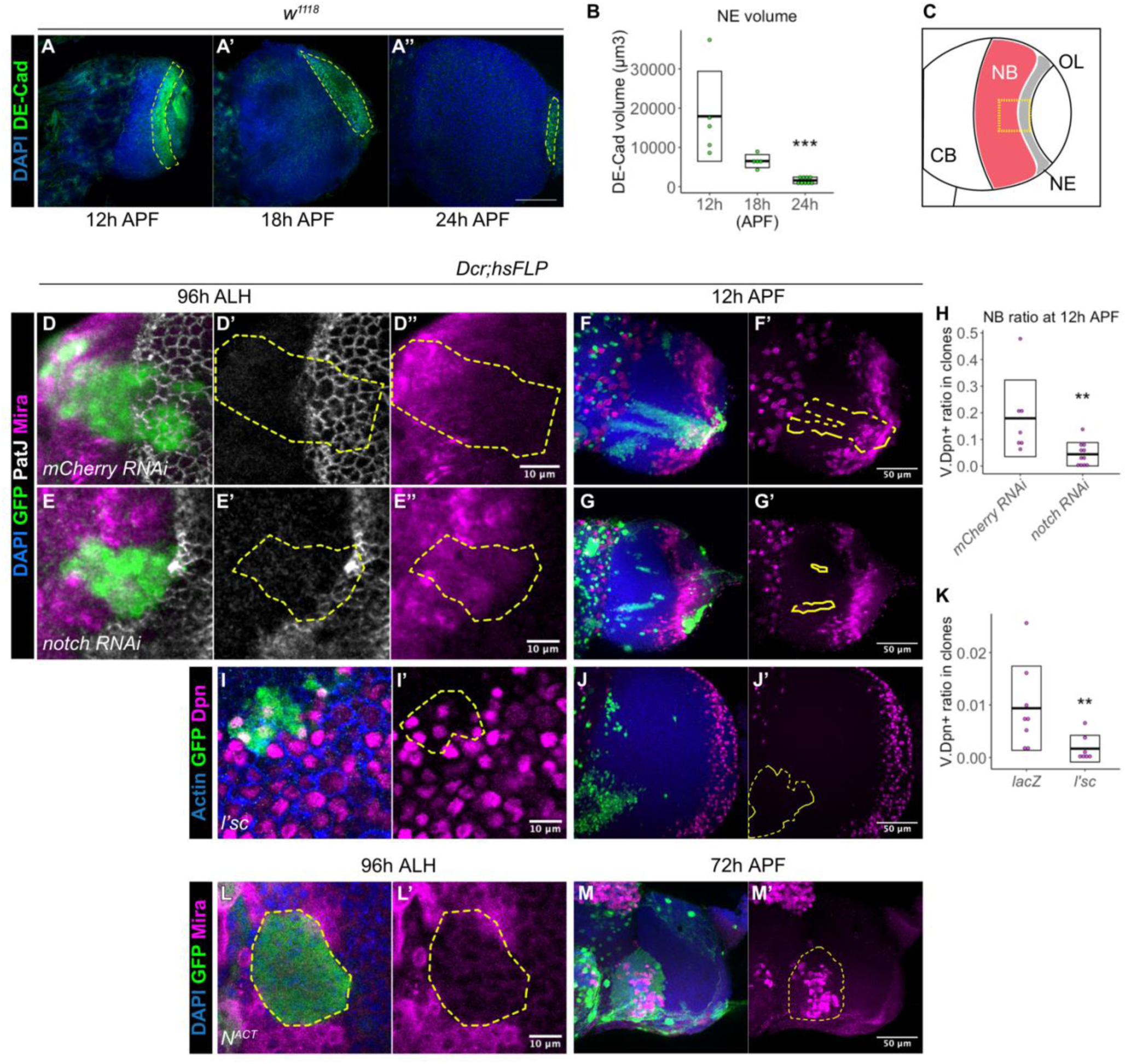
The termination of medulla NBs is scheduled by the timing of the larval NE-NB transition. **(A-A’’)** The NE pool marked by DE-Cad (green, dashed lines) becomes gradually depleted from 12 hr to 24 hr APF. DAPI (blue) **(B)** Quantification of DE-Cad volumes at 12 hr, 18 hr, and 24 hr APF. Kruskal-Wallis test and Dunn’s test to correct for multiple comparisons: ***p = 0.0003. 12 hr: n = 5, m = 17922 ± 5129. 18 hr: n = 5, m = 6508 ± 747.0. 24 hr: n = 9, m = 1597 ± 280.9. **(C)** Schematic depicting the larval OL at 120 hr ALH, where the NE-NB transition is boxed, and is the region of interest in (D-M). **(D-E’)** Representative images of *hsFLP* clones (dashed lines) at 120 hr ALH, induced with (D-D’’) *UAS-mCherry RNAi* and (E-E’’) *UAS-notch RNAi*. GFP (green), PatJ (grey), Mira (magenta). **(F-G’)** Representative maximum projections of the OL at 12 hr APF, in which *hsFLP* clones (dashed lines) are induced with (F-F’) *UAS-mCherry RNAi* and (G-G’) *UAS-notch RNAi*. DAPI (blue), GFP (green), Mira (magenta). **(H)** Quantification of the NB ratio in clones in the OL at 12 hr APF in which *hsFLP* clones are induced with *UAS-mCherry RNAi* and *UAS-notch RNAi*. Mann-Whitney test: **p = 0.0041. *mCherry RNAi*: n = 7, m = 0.179 ± 0.054. *notch RNAi*: n = 11, m = 0.044 ± 0.013. **(I-I’)** Representative *hsFLP* clone (dashed line) at 120 hr ALH, induced with *UAS-l’sc*. **(J-J’)** Representative maximum projections of the OL at 12 hr APF, in which *hsFLP* clones (dashed line) are induced with *UAS-l’sc*. Actin (blue), GFP (green), Dpn (magenta) **(K)** Quantification of the NB ratio in clones in the OL at 12 hr APF in which *hsFLP* clones are induced with *UAS-lacZ* and *UAS-l’sc*. Mann-Whitney test: **p = 0.0057. *lacZ*: n = 8, m = 0.009 ± 0.003. *l’sc*: n = 7, m = 0.002 ± 0.001. **(L-L’)** Representative *hsFLP* clone (dashed line) at 120 hr ALH, induced with *UAS-N^ACT^*. **(M-M’)** Representative maximum projections of the OL at 72 hr APF, in which *hsFLP* clones (dashed lines) are induced *UAS-N^ACT^*. Note that *N^ACT^* clones do not generate NBs until late pupal development. DAPI (blue), GFP (green), Mira (magenta)

To test if the depletion of the NE can change the timing of NB termination, we manipulated Notch signalling, a negative regulator of the NE-NB transition <u>(Egger et al., 2010; Wang et al., 2011; Yasugi et al., 2010)</u>, where Notch signalling promotes NE self-renewal and prevents their differentiation into NBs <u>(Egger et al., 2010; Wang et al., 2011; Yasugi et al., 2010)</u>. Control and *notch* RNAi clones were induced in the medulla NE at 24 hr after larval hatching (ALH) and examined at 96 hr ALH. Control clones spanned the NE and NB regions, demarcated by a sharp NE-NB boundary indicated by PatJ (NE) and Mira (NB) (Figure 2D). In contrast, *notch* RNAi clones consisted mostly of only NBs, suggesting that Notch inhibition has caused an accelerated NE-NB transition (Figure 2E) <u>(Egger et al., 2010; Wang et al., 2011; Yasugi et al., 2010)</u>. By 12 hr APF, we found that *notch* RNAi clones were smaller in size and contained very few NBs compared to control (Figure 2F-H). Thus, it is likely that an accelerated NE-NB transition caused by *notch* RNAi resulted in a premature depletion of the NE which in turn led to the early termination of NBs. To test this further, we overexpressed the proneural gene *lethal of scute* (*l’sc*), which has also been shown to promote NE-NB transition during the larval stages <u>(Yasugi et al., 2008)</u> (Figure 2I). Similar to *notch* RNAi clones, *l’sc* overexpression caused medulla NBs to undergo premature termination at 12 hr APF (Figure 2J-K). Together, our data indicate that the premature depletion of the NE results in earlier medulla NB termination.

Conversely, we found that the overexpression clones of Notch signalling via *N^ACT^* caused a delay in the NE-NB transition, resulting in clones consisting only of NE cells (and no NBs) at 96 hr ALH (Figure 2L). We only started to observe NBs in *N^ACT^* clones in the OLs (Figure 2M, n = 8/8 lobes) at 72 hr APF, a time point where control medulla NBs have long disappeared. These medulla NBs continued to persist into the adult OL (data not shown). Together, our results suggest that early depletion of the NE results in the early cessation of medulla NBs, whereas delayed formation of the NB from the NE results in late termination of medulla NBs. Thus, the timing of the NE-NB transition determines when medulla NBs terminate.

### Medulla NBs cease divisions via cell death

Next, we examined how medulla NBs undergo termination. First, we examined mechanisms that are responsible for the termination of other NB lineages. Mushroom Body NBs are the last NBs to undergo termination <u>(Ito & Hotta, 1992)</u>. These NBs are eliminated via a combination of apoptosis and autophagy (Siegrist *et al*, 2010). To test if medulla NBs terminate via similar mechanisms, we first assessed cell death, where we assayed for the expression of the apoptotic marker Caspase-1 (Dcp-1) in the medulla NBs marked by Dpn::GFP. At 12 hr APF, we observed that some NBs expressed Dcp-1 (Figure 3A). To test if apoptosis is required for the termination of medulla NBs, we inhibited cell death via the overexpression of the baculovirus anti-apoptotic gene *p35* in clones as well as in medulla NBs using *eyR16F10-GAL4*. Upon the inhibition of cell death, we observed a small number of persistent NBs at 24 hr APF through both approaches (Figure 3B-D, Supplementary Figure 2A-B, E). However, no persistent NBs were found at 48 hr APF, suggesting that additional factors may be involved to terminate medulla NBs (Supplementary Figure 2D, n = 5/5 lobes).

**Figure 3:**
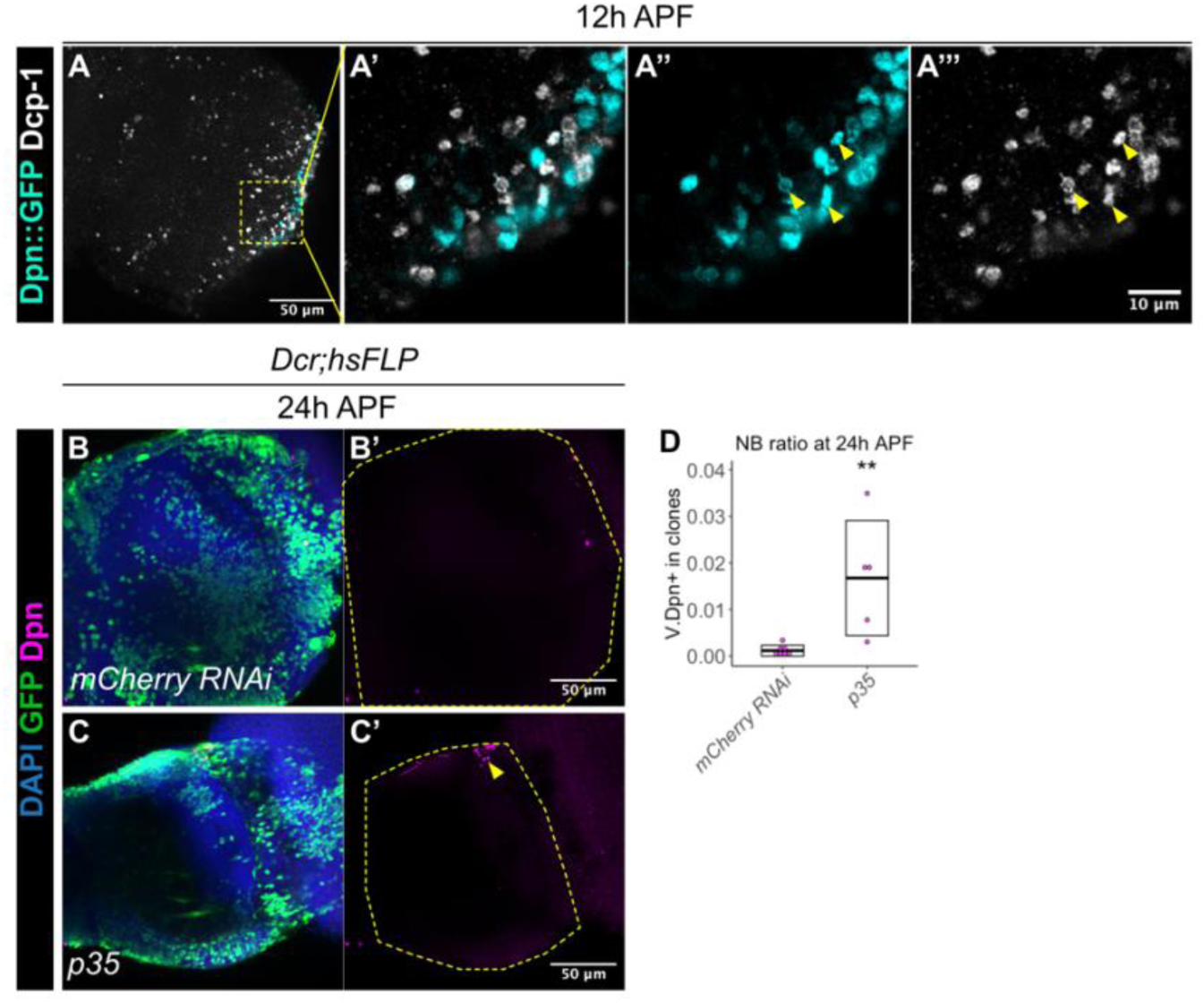
A subset of medulla NBs terminate via cell death. **(A)** At 12 hr APF, a subset of medulla NBs marked by Dpn::GFP (cyan) express the apoptotic marker Dcp-1 (grey) (arrowheads). A’-A’’’ is the magnified view of the boxed inset in A. **(B-C’)** Representative maximum projections of the OL (dashed line) at 24 hr APF with *hsFLP* clones induced with (B-B’) *UAS-mCherry RNAi* and (C-C’) *UAS-p35*. Ectopic NBs are indicated by arrowhead. DAPI (blue), GFP (green), Dpn (magenta). The same *hsFLP>mCherry RNAi* representative image is used in Supplementary figure 1F and Figure 4E. **(D)** Quantification of the NB ratio in *hsFLP* induced with *UAS-mCherry RNAi* and *UAS-p35* at 24 hr APF in the OL. Mann-Whitney test: **p = 0.005. *mCherry RNAi*: n = 7, m = 0.001 ± 0.0004. *p35*: n = 5, m = 0.017 ± 0.006.

Next, we asked if autophagy is required to eliminate medulla NBs. To address this, we inhibited autophagy via the expression of a RNAi against Atg1 in the medulla NBs with *eyR16F10-GAL4* <u>(Xu *et al*, 2015)</u>. We found that the timing of medulla NB termination was not significantly affected by the inhibition of autophagy (Supplementary Figure 2C, E). Together, our data suggest that cell death, and not autophagy, makes a minor contribution to medulla NB termination.

### Medulla NBs undergo Pros-mediated termination

In type I NBs of the VNC, the nuclear localisation of a differentiation factor called Prospero (Pros) precedes NB termination via size-symmetric cell division <u>(Choksi et al., 2006; Li et al., 2013; Maurange et al., 2008</u>). To assess if medulla NBs also undergo Pros-mediated terminal differentiation, we first examined Pros expression together with the pan-NB reporter Dpn::GFP at 12 hr APF. We found that many Dpn^+^ NBs expressed nuclear Pros (Figure 4A). Then, we examined the expressions of Dpn::GFP with Miranda (Mira) marks mature NBs and asymmetric cell division at 12 hr APF. Interestingly, we occasionally observed some small Dpn^+^ Mira^-^ cells, that form doublets (Figure 4B), which we think are likely to be progenies of NB symmetric divisions. Therefore, it is possible that medulla NBs also undergo size symmetric divisions via Pros localisation like NBs of other lineages. To test if Pros is required for the termination of medulla NBs, we induced *pros* RNAi clones (Shaw *et al*, 2018). This manipulation caused the formation of ectopic NBs in the deep layers of the medulla where neurons reside during late larval stages (Figure 4C-B). In the *pros RNAi* clones, many persistent NBs were found at 24 hr APF in contrast to the control (Figure 4E-G). Together, our data suggest Pros is required to promote differentiation of the medulla NBs, and consequently shut down medulla neurogenesis during pupal life (Figure 4H).

**Figure 4:**
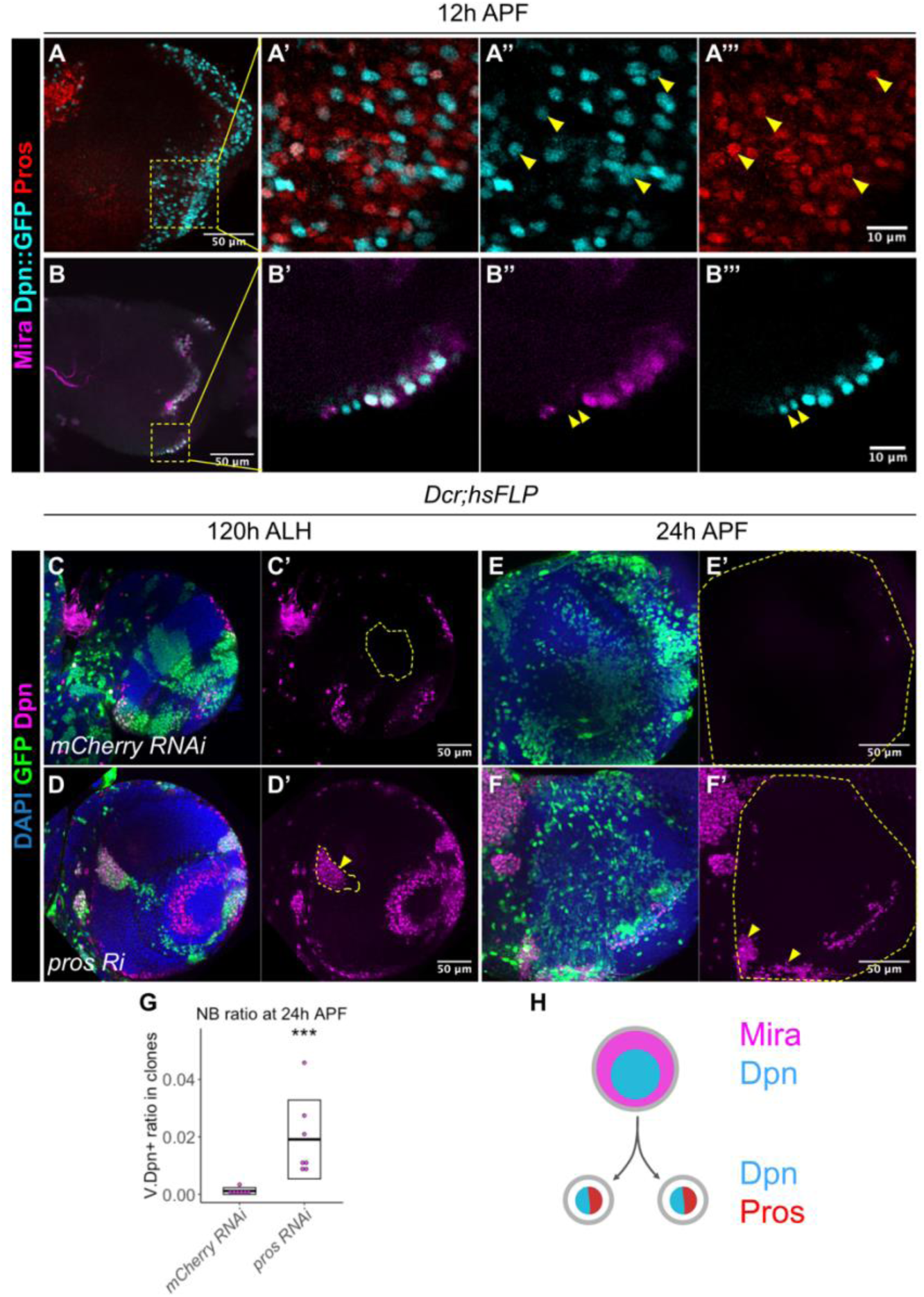
The termination of medulla NBs requires Pros-mediated size symmetric division. **(A-A”’)** Representative images of the OL at 12 hr APF. Some NBs marked by Dpn::GFP (cyan) in the medulla co-express Pros (red) (arrowheads). A’-A’’’ are magnified images of boxed inset in A. **(B-B’’’)** Representative images of the OL at 12 hr APF. On the superficial layer where medulla NBs reside, we have occasionally observed a pair of similarly-sized cells at the most medial position (arrowheads), that express Dpn::GFP (cyan) but not Mira (magenta). B’-B’’’ are magnified images of boxed inset in B. **(C-D’)** Representative images of the deep layers of the medulla with *hsFLP* clones (dashed lines) driving (C-C) *UAS-mCherry RNAi* and (D-D’) *UAS-pros RNAi* at 120 hr ALH. Arrowhead indicates ectopic NBs in the deep section of the medulla. **(E-F’)** Representative maximum projects of an OL (dashed line) with *hsFLP* clones driving (E-E’) *UAS-mCherry RNAi* and (F-F’) *UAS-pros RNAi* at 24 hr ALH. Arrowhead indicates persistent NBs. The same *hsFLP>mCherry RNAi* representative image is used in Supplementary figure 1F and Figure 3B. DAPI (blue), GFP (green), Dpn (magenta). **(G)** Quantification of the NB ratio in clones in the OL at 24 hr APF, in which *hsFLP* clones are induced with *UAS-mCherry RNAi* and *UAS-pros RNAi*. Mann-Whitney test: ***p = 0.0006. *mCherry RNAi*: n = 7, m = 0.001 ± 0.0004. *pros RNAi*: n = 7, m = 0.019 ± 0.005. **(H)** Schematic depicting a medulla NB expresses nuclear Pros to promote symmetric division.

### Tailless is involved in controlling the timing of medulla NB termination

In the OL, medulla NBs sequentially express an array of temporal transcription factors as they age to promote cellular diversity <u>(Konstantinides et al., 2022; Li et al., 2013; Zhu et al., 2022)</u>. The simplified temporal series includes Homothorax (Hth), Eyeless (Ey), Sloppy-paired (Slp), Dichaete (D) and Tailless (Tll) (Figure 5A). It was previously shown that nuclear Pros localisation occurred during the Tll^+^ temporal window <u>(Li et al., 2013)</u>; and that the glial regulatory gene, Gcm, is expressed in the Tll^+^ NBs prior a switch from neurogenesis to gliogenesis <u>(Konstantinides et al., 2022; Li et al., 2013; Zhu et al., 2022)</u>.

**Figure 5:**
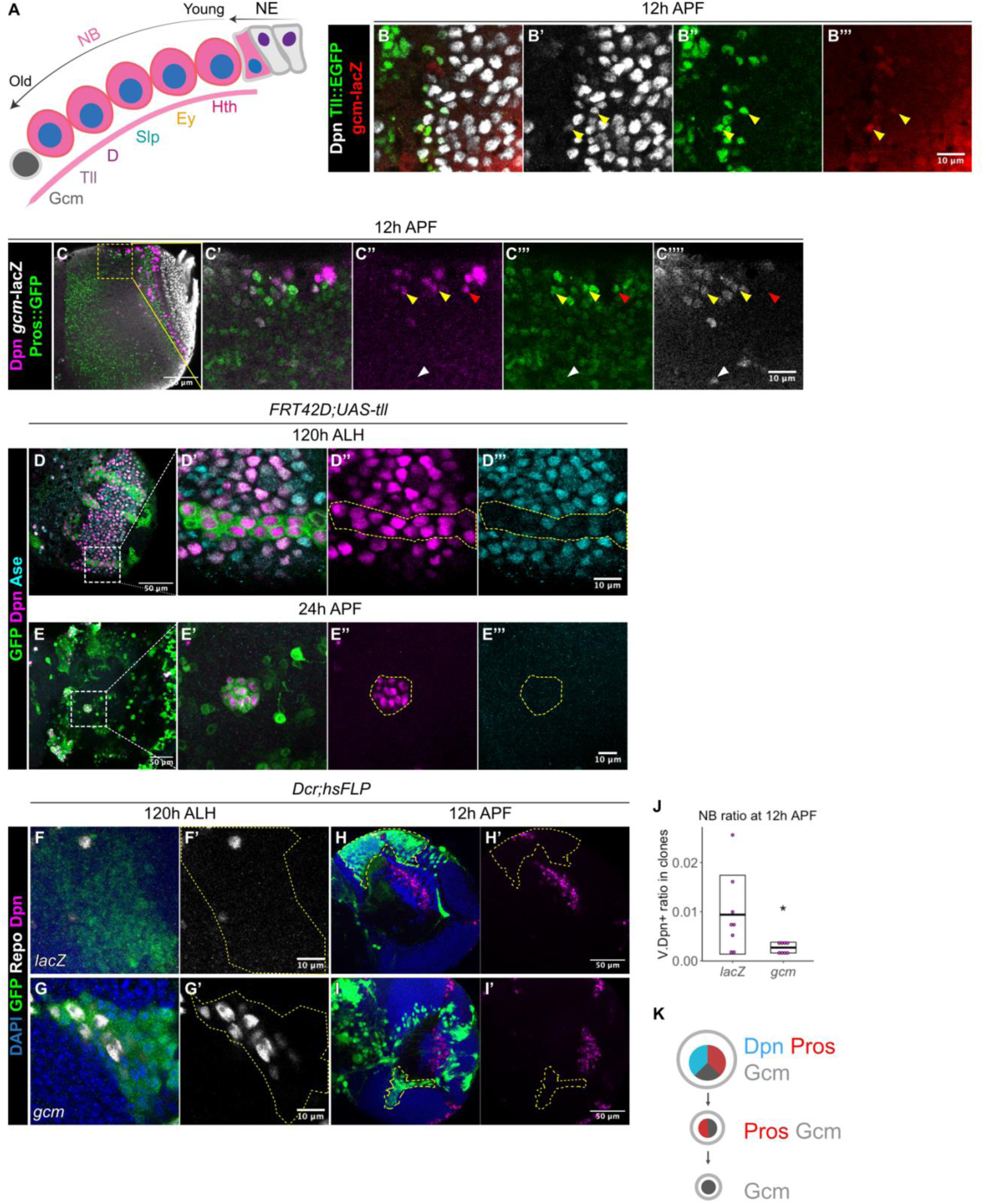
Gcm is sufficient to induce gliogenesis in the medulla at the expense of NBs. **(A)** Schematic of the temporal progression in the OL medulla. **(B-B’’’)** Representative images of the superficial layer of the medulla NBs at 12 hr APF. Some of the most medial NBs (arrowheads) express Dpn (gray), Tll::EGFP (green), and *gcm*-lacZ (red). **(C-C’’’’)** Representative images of the OL at 12 hr APF. Here, some of the most medial NBs are Dpn^+^Pros^+^gcm^+^ (yellow arrowheads) whereas some are Dpn^+^Pros^+^gcm^-^ (red arrowheads) on the superficial layers. In the deep layers, some cells are Dpn^-^Pros^-^gcm^+^ (white arrowheads). C’-C’’’ are magnified images of C. **(D-D’’’)** Representative images of the OL at 120 hr ALH, in which MARCM clones overexpressed for Tll are induced. Box indicates inset shown in L’-L’’’. In the clone (dashed line), medulla NBs expressed low levels of Ase compared to wildtype medulla NBs outside of the clone. D’-D’’’ are magnified images of D. **(E-E’’’)** Representative maximum projections of the OL at 24 hr APF, in which MARCM overexpressed for Tll are induced. In the clone (dashed line), ectopic NBs do not express Ase. E’-E’’’ are magnified images of E. GFP (green), Dpn (magenta), Ase (cyan). **(F-G’)** Representative *hsFLP* clones (dashed lines) induced with (F-F’) *UAS-lacZ*, and (G-G’) *UAS-gcm* at 120 hr APF. **(H-I’)** Representative images of the OLs (dashed lines) at 12 hr APF, in which *hsFLP* clones are induced with (H-H’) *UAS-lacZ*, and (I-I’) *UAS-gcm*. **(J)** Quantification of the NB ratio in clones in the OLs at 12 hr APF, in which *hsFLP* clones are induced with *UAS-lacZ* and *UAS-gcm*. Mann-Whitney test: *p = 0.0379. *lacZ*: n = 8, m = 0.009 ± 0.003. *gcm*: n= 8, m = 0.003 ± 0.0004. **(K)** Schematic depicting a Dpn^+^ medulla NB expresses nuclear Pros and the glial cell fate determinant Gcm to induce a switch from neurogenic-to-gliogenic program.

At 12 hr APF, we found some Tll^+^ NBs were also *gcm*^+^ (*gcm*-lacZ, Vincent *et al*, 1996) (Figure 5B), indicating that a neurogenic to gliogenic switch could also contribute towards the disappearance of NBs at this time point. Consistent with this, at 12 hr APF, we observed some superficially located Dpn^+^ NBs that expressed both the differentiation marker Pros and *gcm*-lacZ (Figure 5C, yellow arrowheads), suggesting that these NBs could be undergoing a NB to glioblast cell fate transition. Deeper within the medulla, we found *gcm*^+^ cells which express neither Dpn nor Pros (Figure 5C, white arrowhead), that are likely to be the progenies of the glioblasts. Finally, we found a few medially located Dpn^+^ NBs that expressed Pros but not *gcm*-lacZ (Figure 5G, red arrowhead). This is consistent with what we observed in Figure 4A-B, where some NBs undergo terminal differentiation through likely via symmetric cell division.

To test if members of the temporal transcription factor series can alter NB persistence and prevent the neurogenic to gliogenic switch, we first examined the role of Tll, which defines the temporal window where terminal differentiation as well as the gliogenic switch occur <u>(Li et al., 2013; Zhu et al., 2022</u>). First, we induced *tll* overexpression clones and found that *tll* overexpression caused the formation of ectopic NBs in the larval CNS (Figure 5D, and these NBs persisted at 24h APF (Figure 5E, n = 4/4 lobes). *tll* overexpression has been previously shown to convert type I NBs to a type II NB cell fate that are capable of tumour formation in type I NBs of the CB <u>(Hakes & Brand, 2020)</u>. Indeed, we found that the *tll* overexpressing NBs were mostly Dpn^+^Ase^-^ which is characteristic of type II NBs (Figure 5D-E). Therefore, it is possible that *tll* overexpression caused type I to type II conversions that accounts for why these clones failed to undergo termination on time.

Consistent with reports that Tll is not required for glial production in the larval medulla NBs <u>(Zhu et al., 2022)</u>, we found that inhibition of *tll* via a validated RNAi <u>(Hakes & Brand, 2020)</u> was not able to alter the number of glial cells in the larval medulla (Supplementary Figure 3A, B, E), nor the timing of medulla NB termination in pupal stages (Supplementary Figure 3C, D, F). Moreover, the knockdown of *tll* did not alter the number of NBs undergoing cell death assayed via Dcp-1 expression (Supplementary figure 3G-H) nor the expression of Pros which promotes terminal size symmetric divisions (Supplementary figure 3I-J). Together, while Tll is not required for medulla NB termination, ectopic expression of Tll can induce persistent NBs. Furthermore, Tll acts independently of cell death and symmetric NB termination mechanisms.

To assess whether early temporal transcription factors are required for the termination of medulla NBs, we overexpressed Hth and inhibited Ey or Scarecrow (Scro) <u>(Konstantinides et al., 2022; Zhu et al., 2022</u>) via RNAi in clones, which prevents the progression through the temporal series. Nevertheless, none of these manipulations influenced the timing of NB termination (data not shown). Taken together, altering temporal progression is not sufficient to affect the timing of medulla NB termination in the pupal stages.

### Gcm-induced gliogenic switch can affect the timing of NB termination

We next assessed if manipulating the gliogenic switch can affect the timing of medulla NB termination. Here, we induced *gcm* overexpression clones using a reagent previously validated in <u>Zhu et al., 2022</u>. This manipulation was able to induce the formation of ectopic glial cells during larval neurogenesis (Figure 5F-G). At 12 hr APF, we recovered significantly fewer NBs in the OL upon *gcm* overexpression (Figure 5H-J), suggesting that the induction of a gliogenic switch can cause precocious medulla NB termination. To assess if the downregulation of *gcm* can cause prolonged NB persistence in the medulla, we knocked down *gcm* using a previously validated RNAi <u>(Zhu *et al*, 2022)</u> in clones. This manipulation was able to inhibit the expression of the glial cell marker Repo in clones during larval stages (Supplementary figure 3K-L). However, no persistent medulla NBs was found at 24 hr APF (Supplementary figure 3M-N, n = 5/5 lobes). This suggests that while the downregulation of *gcm* is not sufficient to induce NB persistence, *gcm* overexpression can cause a neurogenic to gliogenic switch and thereby, premature termination of NBs.

## Discussion

Robust brain development requires the tight coordination between stem cell maintenance and differentiation. In the *Drosophila* OL, there are two main stem cell pools, the NE cells that divide symmetrically, and the NBs. Medulla NBs are derived from the OPC NE via the proneural wave, which switch to asymmetric division where they self-renew and give rise to GMCs that will produce post-mitotic neurons and glial cells of the medulla cortex <u>(Egger et al., 2007; Li et al., 2013; Yasugi et al., 2008)</u>. Our study shows that during early pupal development, medulla NBs terminate via a combination of mechanisms including cell death, Pros-mediated differentiation and a switch from neurogenesis to gliogenesis. While factors such as ecdysone signalling and OxPhos of the NBs are involved in NB termination in other lineages outside of the OL <u>(Homem et al., 2014; Syed et al., 2017; van den Ameele & Brand</u>, <u>2019; Yang et al., 2017)</u>, we found that these factors do not play a significant role in the termination of medulla NBs. Instead, the main regulator of the timely NB cessation in the pupal medulla is regulated by the timing of the conversion of NE-NB conversion during larval stages. Intriguingly, while a specific cascade of temporal identity transcription factors in the medulla NBs has been shown to generate the cellular diversity of medulla neurons and glia <u>(Konstantinides et al., 2022; Li et al., 2013; Zhu et al., 2022)</u>, temporal progression in the medulla NBs appears dispensable for the timing of NB cessation in the pupal stages (Figure 6). Diseases such as microcephaly are mediated by the premature depletion of the NE pool, or through a precocious commitment to neurogenic divisions <u>(Götz & Huttner, 2005)</u>. Consistent with this, our data present evidence that during development, the regulation of the NE stem cell pool, and its timely conversion into NSCs is of critical importance to the overall size and function of the brain.

**Figure 6:**
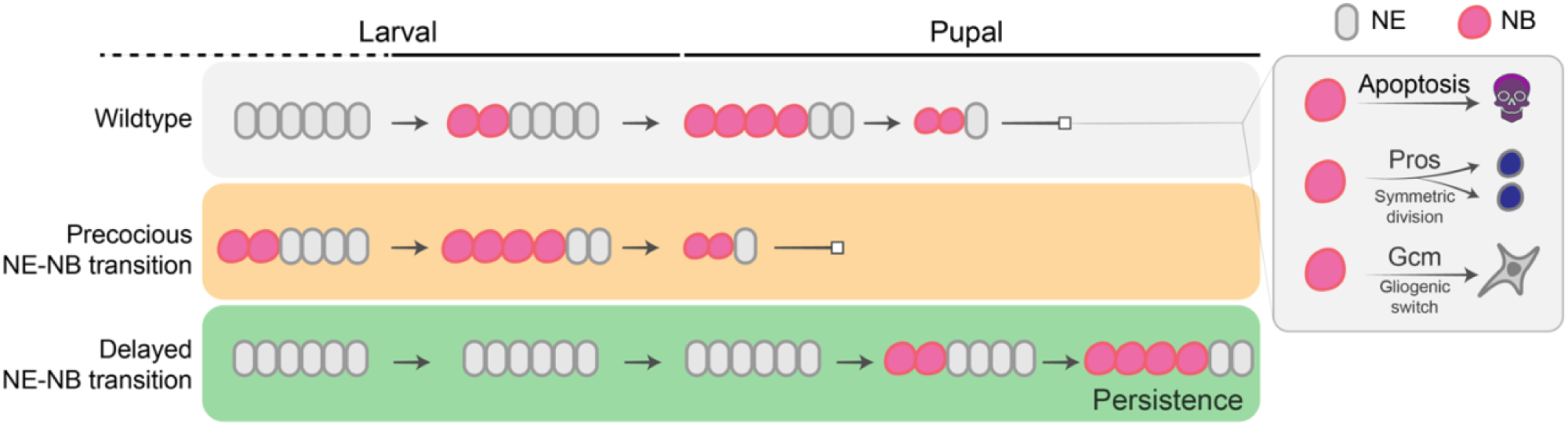
Working model of the termination of medulla NBs. Timing of NB termination in the early pupal stages is scheduled by the timing of the NE-NB transition in the larval stages. Medulla NBs can be terminated via three mechanisms: (1) cell death, (2) Pros-mediated symmetric division and (3) Gcm-mediated gliogenic switch of the NBs.

### NE-NB transition determines the timely termination of medulla NBs

Our data support a model whereby once the NE pool is exhausted, no new NBs can be generated. NBs transit through their temporal series to terminally differentiate to become glial cells or neurons, which marks the end of medulla neurogenesis. Thus, unlike type I NB lineages in which NB termination is scheduled by ecdysone signalling-mediated temporal progression at the level of the NBs <u>(Syed et al., 2017; Yang et al., 2017)</u>, in the OL, NB termination is pre-determined in the NE. In the OL, besides medulla NBs, NE cells also sequentially express an array of temporal factors, such as Chinmo and Broad downstream of ecdysone signalling <u>(Arain et al., 2022; Dillard et al., 2018; Lanet et al., 2013; Zhou et al., 2019)</u>. Moreover, inhibiting the temporal Chinmo-to-Broad expression switch in the NE was shown to delay the NE-NB transition, resulting in tumours with persistent NBs in the adult OLs (<u>Dillard et al., 2018; Zhou et al., 2019</u>). The differences in the regulation of NB termination in the OL medulla versus other lineages may be due to some fundamental differences in the origins of these NBs. For example, in the OL, NBs are derived from the NE, that undergoes rapid growth during larval development <u>(Egger et al., 2007; Lanet et al.</u>, <u>2013; Yasugi et al., 2008)</u>. Therefore, NB termination in the OL is scheduled at the level of the NE, likely by the activation of ecdysone signalling during late larval development. In contrast, NBs from other lineages are produced directly from the neuroectoderm in the embryo without undergoing the transition from the NE <u>(Harding & White, 2018)</u>. Hence, their termination mechanism acts predominantly in the NBs during pupal development.

### Cell death makes a small contribution to NB termination

Consistent with previous reports, we found that cell death occurred in a subset of medulla NBs at 12 hr APF <u>(Hara *et al*, 2018</u>). However, inhibition of cell death did not fully prevent medulla NBs from termination in the pharate OLs (Supplementary Figure 2D), suggesting that NBs can adopt alternative routes for timely termination before adulthood. From our data, it is not yet clear whether a specific subset of medulla NBs undergo cell death. Although the timing of cell death in the medulla NBs coincides with a small pulse of ecdysone around 10 hr APF <u>(Riddiford, 1993)</u>, we showed that inhibition of ecdysone signalling in the medulla NBs did not phenocopy the inhibition of apoptosis (Supplementary Figure 1B-E). Therefore, unlike neurons in the OL which require ecdysone signalling for cell death during pupal development <u>(Hara et al., 2013)</u>, medulla NBs undergo cell death independently of ecdysone signalling. Altogether, while some NBs terminate proliferation through cell death, it is likely that cell death of the NBs makes a minor contribution to the termination of neurogenesis in the medulla.

### A neurogenic-gliogenic switch and symmetric divisions cause NB cessation

Our data show that many medulla NBs displayed nuclear Pros expression during early pupal stages (Figure 3A). Consistent with the essential role of Pros in promoting differentiation <u>(Choksi et al., 2006; Maurange et al., 2008)</u>, *pros* knockdown caused the persistence of the NBs in the pupal OLs (Figure 3C-G), similar to other regions of the brain. In addition to differentiation, another mechanism responsible for the cessation of neurogenesis, is the commencement of gliogenesis. We found glioblasts in the early pupal medulla, which make glial cells at the end of the temporal transcription factor series, express the NB marker Dpn, the glial marker Gcm as well as the differentiation marker Pros (Figure 5B-C). As such, it is possible that similar to the embryonic glioblasts, Pros is required for glial cell fate acquisition via Gcm <u>(Freeman & Doe, 2001; Ragone et al., 2001)</u>. As Gcm knockdown did not induce any persistent medulla NBs during pupal stages (Supplementary Figure 3M-N), the presence of Pros may likely induce symmetric division of the NBs, resulting in their termination.

Together, cell death, symmetric cell division and a gliogenic switch appear to be employed as fail-safe mechanisms that help to eliminate the NB progenitor pool in the pupal OL. Hence, it would be interesting to explore in the future the cross-regulatory relationship between these termination programs.

## Materials and Methods

### Fly husbandry

Fly stocks were reared on standard media at 25°C. For larval dissection, brains were dissected at wandering L3 stages (120 hr ALH). For pupal dissection, white pupae were selected and allow to age until the desired stages (12 hr, 24 hr, 30 hr, 48 hr, and 72 hr APF).

Wildtype fly was *w^1118^*. *eyR16F10-GAL4* (BL 48737) was used to induce expression of transgenes in a subset of medulla NBs and their progeny from larval development <u>(Veen *e*</u>*t <u>al</u>*<u>, 2023)</u>. Knockdown and overexpression clones were induced by flip-out system using *hsFLP,act>CD2>GAL4;UAS-Dcr2,UAS-GFP* (a gift from Lei Zhang). Flip-out clones were induced at 24 hr ALH for 10 min. *FRT42D;UAS-tll* clones were induced using the MARCM system: *UAS-mCD8::GFP,hsFLP;FRT42D, tub-GAL80; tub-GAL4* (a gift from Alex Gould). Clones were induced at 48 hr ALH for 15 min.

The fly strains used were as followed: *Dpn::GFP* (Morin *et al*, 2001), *FRT42D* (a gift from Alex Gould), *gcm-lacZ* (BL 5445), *Tll::EGFP* (BL 30874), *Pros::GFP* (VDRC 318463), *UAS-Atg1 RNAi* (BL 26731), *UAS-dp110^CAAX^*(BL 25908), *UAS-EcR RNAi* (VDRC 37058), *UAS-EcR^DN^* (BL 9451), *UAS-gcm* (BL 5446), *UAS-gcm RNAi* (VDRC 110539), *UAS-l’sc* (BL 51670), *UAS-lacZ* (BL 8529), *UAS-mCherry RNAi* (BL 35785), *UAS-med27 RNAi* (BL 34576), *UAS-N^ACT^* (a gift from Helena Richardson), *UAS-ND75 TRiP* (BL 33911), *UAS-notch RNAi* (VDRC 100002), *UAS-p35* (BL 5072), *UAS-pros RNAi* (BL 26745), *UAS-tll* (a gift from Claude Desplan), *UAS-tll RNAi* (VDRC 330031), *UAS-D* (BL 8861).

### Immunostaining

Larval and pupal brains were dissected out in phosphate buffered saline (PBS), fixed in 4% formaldehyde for 20 min and rinsed three time in 0.5% PBST (PBS + 0.5% Triton) at room temperature. For immunostaining, brains were incubated in primary antibodies overnight at 4°C, followed by two washes in 0.5% PBST and then an overnight secondary antibody incubation at 4°C or 3 hrs at room temperature. Brains were afterwards rinsed two times in 0.5% PBST and incubated in 50% glycerol in PBS for 20 min. Samples were mounted in 80% glycerol in PBS, on glass slides and sealed with coverslips (22×22 mm, No.1.5, Knittel) for image acquisition. The primary antibodies used were: rat anti-Dpn (1:200, Abcam 195172), chick anti-GFP (1:1000, Abcam 13970), mouse anti-EcR^common^ (1:50, DSHB Ag10.2), rabbit anti-Dcp-1 (1:100, Cell Signalling 95785), rat anti-Mira (1:100, Abcam 197788), rat anti-Pros (a gift from Fumio Matsuzaki), rabbit anti-β-Galactosidase (1:200, a gift from Helena Richardson), rabbit anti-Ase (a gift from Lily Jan and Yuh Nung Jan), mouse anti-Repo (1:50, DSHB 8D12), and rat anti-DE-Cad (1:200, DSHB DCAD2), rabbit anti-PatJ (1:500, a gift from Helena Richardson). Secondary donkey antibodies conjugated to Alexa 555 and Alexa 647, and goat antibodies conjugated to Alexa 405, 488, 555, and 647 (Molecular Probes) were used at 1:500.

### Imaging and image processing

Images were acquired using Olympus FV3000 confocal microscope with 40x (NA 0.95, UPLSAPO) and 60x (NA 1.30, UPLSAPO) objectives. Δz = 1.5 μm. Images were processed using Fiji (https://imagej.net/Fiji). Single section or maximum projections were shown as representative images in the figures.

Quantification was performed in Fiji or via 3D reconstruction in Imaris (Bitplane). For quantifications of NB numbers (Figures 1C, Supplementary Figure 1E, 2E), number of Dpn^+^ NBs in the medulla/OL were measured. For quantifications of nuclear sizes of medulla NBs (Figure 1E), nuclear diameters identified by Dpn were measured. For quantifications of the ratio of NBs in clones (Figures 2H, 2K, 3D, 4E, 5L, Supplementary figure 1J, 3H), GFP^+^ clones were first detected, and clonal volumes were measured. The GFP^+^ areas were then utilised to make a mask for GFP^+^Dpn^+^ or GFP^+^Mira^+^ NBs within the clones, which was in turn used to measure the total Dpn volume. The NB ratio in clones is calculated as the ratio of Dpn^+^GFP^+^ or GFP^+^Mira^+^ volume over GFP^+^ volume. For quantification of the numbers of glia in the medulla (Supplementary Figure 3G), the total numbers of Repo^+^ cells in the deep section of the medulla were measured in each lobe. For quantification of the NE volumes (Figure 2B), DE-Cad^+^ volumes were measured in each lobe. Images were assembled in Adobe InDesign 2024 and Affinity Publisher 2. Scale bars = 10 μm or 50 μm per indicated in figures.

### Statistical analyses

At least two animals per genotype were used for all experiments. Statistical analyses were performed in GraphPad Prism 9. In graphs, data (crossbars) = mean ± standard error of the mean (SEM). n = lobes unless specified. Graphs were plotted using RStudio. For comparisons between two conditions, P-values were calculated by non-parametric Mann-Whitney tests when data show non-normally distribution. For comparisons between more than two conditions, P-values were calculated by either ordinary one-way ANOVA tests for normally distributed data, or Kruskal-Wallis tests for non-normally distributed data. Dunnet or Dunn’s tests were used to correct for multiple comparisons following one-way ANOVA and Kruskal-Wallis tests, respectively. *(*) p<0.05 (**) p< 0.01, (***) p< 0.001, (****) p < 0.0001, (ns) p>0.05*

## Acknowledgements

We are grateful to Lei Zhang, Alex Gould, Claude Desplan, Fumio Matsuzaki, Helena Richardson, Yuh Nung Jan for generous sharing of fly stocks and antibodies. We would also like to thank Holger Apitz, Édel Alvarez-Ochoa, and Qian Dong for fruitful discussions and critical reading of the manuscript. We also thank Bloomington Drosophila Stock Centre, Vienna Drosophila Resource Centre, and Developmental Studies Hybridoma Bank for fly stocks and antibodies. We would like to also thank OZDros for *Drosophila* quarantine, Peter MacCallum Cancer Institute CAHM for microscopy assistance. LYC’s laboratory is supported by funding from the NHMRC Ideas Grant (APP2011289), and the Peter MacCallum Cancer Foundation.

