## supplemental data for "Neuroepithelial depletion schedules cessation of neurogenesis in the *Drosophila* optic lobes"

Supplementary Figure 1

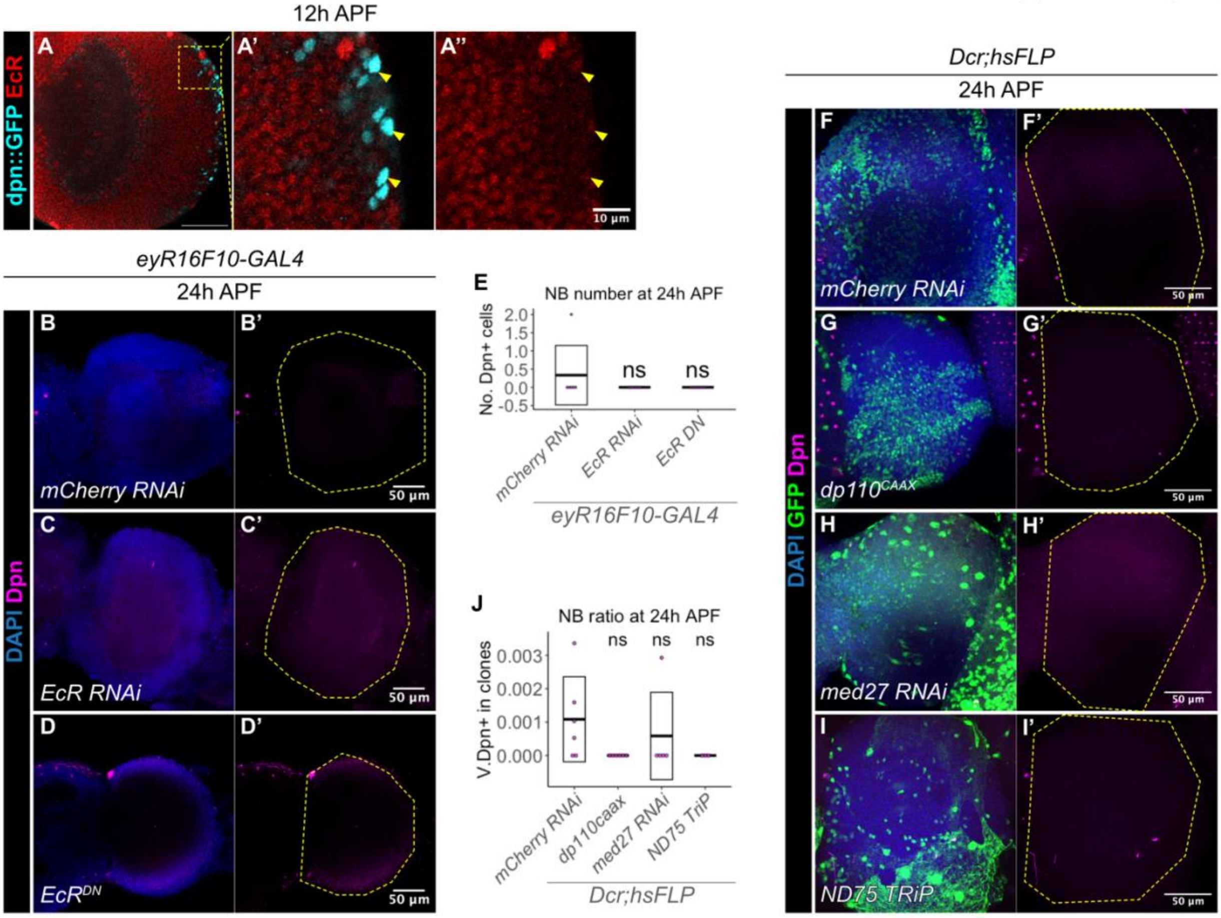


**Figure S1: The termination of medulla NBs does not require ecdysone signalling.**

**(A-A'')** At 12 hr APF, medulla NBs marked by Dpn::GFP (cyan) do not express EcR (red) (arrowheads). A'-A'' is the magnified view of the inset from A.

**(B-D')** Representative maximum projection of the OL at 24 hr APF (dashed line), in which (B-B') *UAS-mCherry RNAi*, (C-C') *UAS- EcR RNAi*, or (D-D') *UAS-EcR$^{DN}$* are driven by *eyR16F10-GAL4*. The same representative image used in B is used in Supplementary figure 2A. DAPI (blue), Dpn (magenta).

**(E)** Quantification of the number of medulla NBs in *UAS-mCherry RNAi*, *UAS-EcR RNAi*, and *UAS-EcR$^{DN}$* driven by *eyR16F10-GAL4* OLs at 24 hr APF. One-way ANOVA test and Dunnet test to correct for multiple comparisons: $p > 0.05$. *mCherry RNAi*: n = 6, m = 0.033 ± 0.333. *EcR RNAi*: n = 6, m = 0.000 ± 0.000. *EcR$^{DN}$*: n = 8, m = 0.000 ± 0.000.

**(F-I')** Representative maximum projections of the OLs in which *hsFLP* clones are induced with (F-F') *UAS-mCherry RNAi*, (G-G') *UAS-dp110$^{CAAX}$*, (H-H') *UAS-med27 RNAi*, and (I-I') *UAS-ND75 TRiP* at 24 hr APF. The same *hsFLP>mCherry RNAi* representative image is used in Figure 3B and Figure 4E. DAPI (blue), GFP clones (green), Dpn (magenta).

**(J)** Quantification of the NB ratio within *hsFLP* clones induced with *UAS-mCherry RNAi*, *UAS-dp110$^{CAAX}$*, *UAS-med27 RNAi*, *UAS-ND75 TRiP* at 24 hr APF in the OL. One-way ANOVA test and Dunnet test to correct for multiple comparisons. *mCherry RNAi*: n = 6, m =

0.001 ± 0.001. *dp110*$^{CAAX}$: n = 6, m = 0.000 ± 0.000. *med 27 RNAi*: n = 5, m = 0.001 ± 0.001. *ND75 TRiP*: n = 3, m = 0.000 ± 0000.

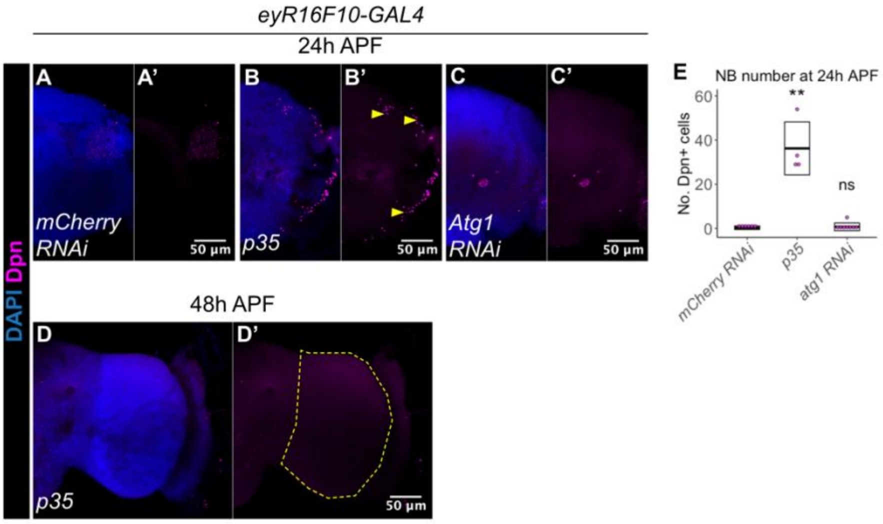


**Figure S2: Autophagy is not necessary for the termination of medulla NBs.**

**(A-C')** Representative maximum projections of the OL in which *eyR16F10-GAL4* drives (A-A') *UAS-mCherry RNAi,* (B-B') *UAS-p35,* (C-C') *UAS-Atg1 RNAi.* (B'B') At 24 hr APF, there are ectopic NBs (arrowheads) in the OL upon the induction of p35. The same representative image used in A is used in Supplementary figure 1B. DAPI (blue), Dpn (magenta).

(D-D') Representative maximum projections of the OL (dashed line) at 48 hr APF, in which *eyR16F10-GAL4* drives *UAS-p35.* DAPI (blue), Dpn (magenta).

**(E)** Quantification of the NB numbers in the OL at 24 hr APF in which *eyR16F10-GAL4* drives *UAS-mCherry RNAi*, *UAS-p35*, and *UAS-Atg1 RNAi.* Kruskal-Wallis test and Dunn's test to correct for multiple comparisons. **p = 0.0041. *mCherry RNAi*: n = 6, m = 0.3333 ± 0.3333. *p35*: n = 4, m = 36.25 ± 6.005. *atg1 RNAi*: n = 8, m = 0.7500 ± 0.6196.

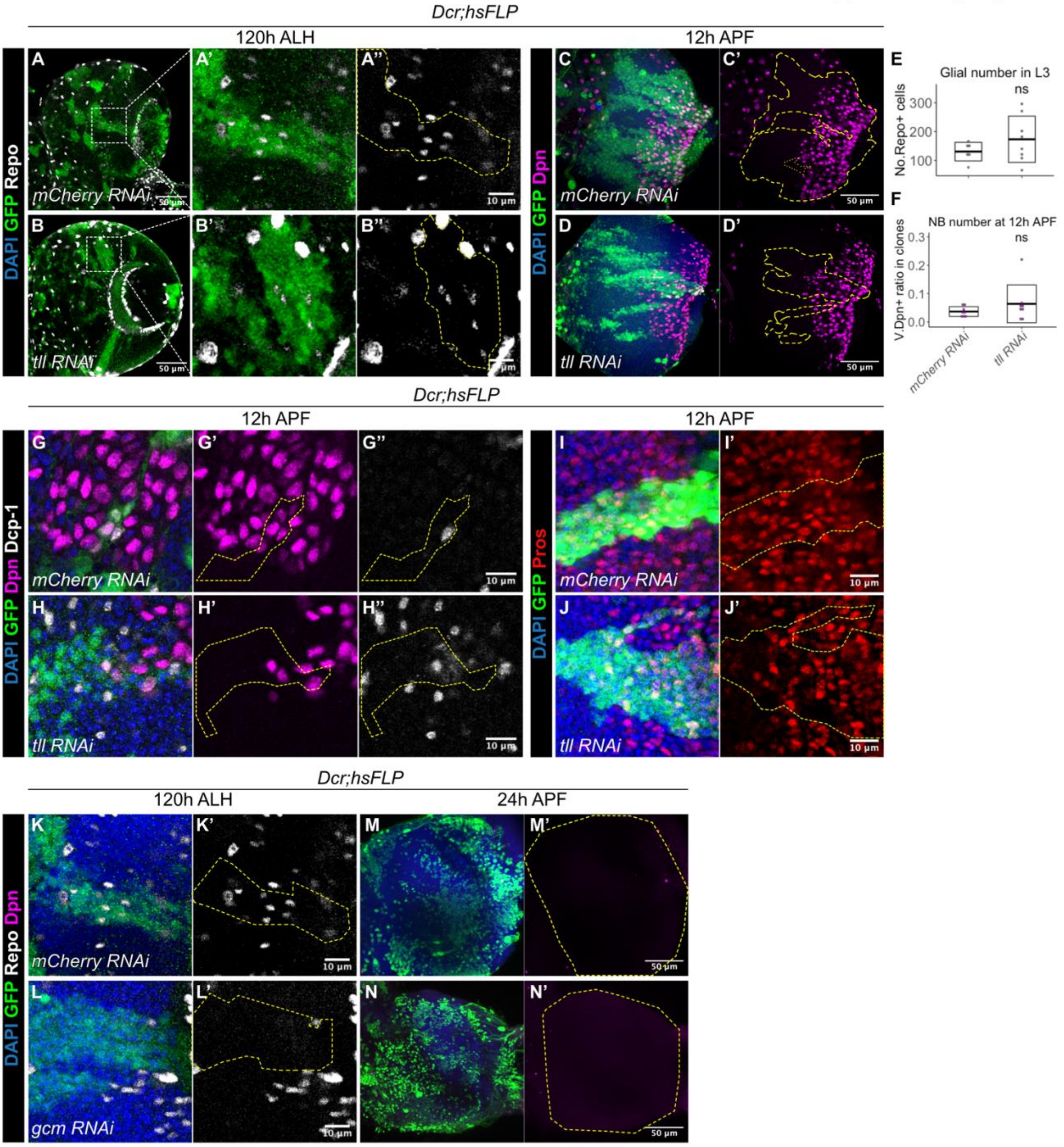


**Figure S3: Tll is not required to schedule the gliogenic switch of the medulla NBs.**

**(A-B'')** Representative images of the deep section of the medulla at 120 hr ALH, in which *hsFLP* clones (dashed lines) are induced with (A-A'') *UAS-mCherry RNAi* and (B-B'') *UAS-tll RNAi*., respectively. GFP (green). Repo (gray). (A'-A'') are magnified images of boxed inset in A, and (B'-B'') are magnified images of boxed inset in B.

**(C-D')** Representative maximum projections of the OLs at 12 hr APF, in which *hsFLP* clones (dashed lines) are induced with (C-C') *UAS-mCherry RNAi* and (D-D') *UAS-tll RNAi*. DAPI (blue), GFP (green), Dpn (magenta).

**(E)** Quantification of the glia number in the medulla in L3 (120 hr ALH), in which *hsFLP* clones are induced with *UAS-mCherry RNAi* and *UAS-tll RNAi*. Mann-Whitney test: $p = 0.4332$. *mCherry RNAi*: $n = 6$, $m = 130.7 \pm 13.22$. *tll RNAi*: $n = 8$, $m = 173.3 \pm 28.39$.

**(F)** Quantification of the NB ratio in clones in the OL at 12 hr APF, in which *hsFLP* clones are induced with *UAS-mCherry RNAi* and *UAS-tll RNAi*. Mann-Whitney test: $p = 0.5054$. *mCherry RNAi*: $n = 8$, $m = 0.037 \pm 0.006$. *tll RNAi*: $n = 8$, $m = 0.064 \pm 0.024$.

**(G-H'')** Representative *hsFLP* clones (dashed lines) at 12hr APF induced with (G-G'') *UAS-mCherry RNAi* and (H-H'') *UAS-tll RNAi*. DAPI (blue), GFP (green), Dpn (magenta), Dcp-1 (grey).

**(I-J')** Representative *hsFLP* clones (dashed line) at 12 hr APF induced with (I-I') *UAS-mCherry RNAi* and (J-J') *UAS-tll RNAi*. DAPI (blue), GFP (green), Pros (red).

**(K-L')** Representative *hsFLP* clones (dashed lines) induced with (K-K') *UAS-mCherry RNAi*, and (L-L') *UAS-gcm RNAi* at 120 hr APF.

**(M-N')** Representative images of the OLs (dashed lines) at 12 hr APF, in which *hsFLP* clones are induced with (M-M') *UAS-mCherry RNAi*, and (N-N') *UAS-gcm RNAi*.
DAPI (blue), Repo (grey), GFP (green), Dpn (magenta).
